# Antisense modulation of IL7R splicing to control sIL7R expression in human CD4^+^ T cells

**DOI:** 10.1101/2022.02.22.481529

**Authors:** Gaddiel Galarza-Muñoz, Debbie Kennedy-Boone, Geraldine Schott, Shelton S. Bradrick, Mariano A. Garcia-Blanco

## Abstract

The interleukin 7 receptor (*IL7R*) is strongly associated with increased risk to develop multiple sclerosis (MS), an autoimmune disease of the central nervous system, and this association is likely driven by upregulation of the soluble isoform of IL7R (sIL7R). Expression of sIL7R is determined by exclusion of the alternative exon 6 from IL7R transcripts, and our previous work revealed that the MS risk allele of the SNP rs6897932 within this exon enhances the expression of sIL7R by promoting exclusion of exon 6. sIL7R potentiates the activity of IL7, leading to enhanced expansion of T cells and increased disability in the Experimental Autoimmune Encephalomyelitis (EAE) murine model of MS. This role in modulating T cell-driven immunity positions sIL7R as an attractive therapeutic target whose expression could be reduced for treatment of MS or increased for treatment of cancers. In this study we identified novel antisense oligonucleotides (ASOs) that effectively control the inclusion (anti-sIL7R ASOs) or exclusion (pro-sIL7R ASOs) of this exon in a dose-dependent fashion. These ASOs provided excellent control of exon 6 splicing and sIL7R secretion in human primary CD4^+^ T cells. Supporting their potential for therapeutic targeting, we showed that lead anti-sIL7R ASOs correct the enhanced exon 6 exclusion imposed by the MS risk allele of rs6897932, whereas lead pro-sIL7R ASOs phenocopy it. The data presented here form the foundation for future pre-clinical studies that will test the therapeutic potential of these ASOs in MS and immuno-oncology.

## INTRODUCTION

The interleukin 7 receptor (*IL7R*) gene is strongly associated with increased risk to develop autoimmune diseases, particularly multiple sclerosis (MS) (Briggs, 2019; Gregory et al., 2007; International Multiple Sclerosis Genetics, 2019; International Multiple Sclerosis Genetics et al., 2007; Lundmark et al., 2007). The *IL7R* gene encodes the IL7R alpha chain (IL7R, CD127), which together with the common cytokine receptor gamma chain (IL2RG, CD132) forms a transmembrane receptor complex for the cytokine interleukin 7 (IL7) in the surface of T cells. IL7 is a pro-survival cytokine and its binding to this receptor complex is essential for the development of T cells in the thymus and the survival of peripheral T cells, and thus the IL7/IL7R axis plays a central role in establishing T cell homeostasis (Carrette and Surh, 2012; Fry and Mackall, 2005; Mazzucchelli and Durum, 2007). In addition to the membrane-bound form of IL7R (hereafter referred to as mIL7R), a secreted isoform named soluble IL7R (sIL7R) can be produced by alternative splicing of IL7R pre-mRNAs. Specifically, sIL7R is formed by exclusion (skipping) of IL7R exon 6, which encodes the single transmembrane span of IL7R (Goodwin et al., 1990). Although a biological function of sIL7R remains elusive, sIL7R has emerged as a pro-immune factor that promotes self-reactive immune responses that underlie autoimmune disorders, with the strongest evidence in MS (Barros et al., 2021). MS is a demyelinating, autoimmune disorder of the central nervous system (CNS), wherein the immune system attacks and destroys the myelin coatings surrounding axons in the brain and spinal cord, leading to progressive neurodegeneration and disability. The precise mechanism(s) underlying this rogue attack on myelin antigens remain unclear; however, it has become increasingly evident that sIL7R plays a role in fueling such pathological attacks in the CNS.

Several lines of evidence support a role of sIL7R as a driver of autoimmunity in MS and other autoimmune disorders. We uncovered the strong and reproducible association of *IL7R* with increased MS risk (Briggs, 2019; Gregory et al., 2007; International Multiple Sclerosis Genetics, 2019; International Multiple Sclerosis Genetics et al., 2007; Lundmark et al., 2007) and proposed this is driven by the effects of the SNP rs6897932 (C/T) within IL7R exon 6 on sIL7R expression. Specifically, we showed the MS risk ‘C’ allele of this SNP increases exclusion of exon 6, thereby enhancing the fraction of IL7R transcripts that encode sIL7R (Gregory et al., 2007). Validating this finding, subsequent studies showed a dose-dependent effect of this SNP on exon 6 splicing and sIL7R secretion, with the highest levels of exon 6 exclusion and circulating sIL7R observed in individuals homozygous for the risk ‘C’ allele, intermediate levels in heterozygous, and the lowest levels in homozygous for the protective ‘T’ allele (Hoe et al., 2010; Lundstrom et al., 2013). Further work from our group revealed that a critical repressor of exon 6 exclusion and sIL7R expression, the RNA helicase DDX39B, is also associated with increased MS risk, which solidifies a role of sIL7R in driving MS (Galarza-Munoz et al., 2017). Importantly, we showed an epistatic interaction between MS risk variants in *IL7R* (rs6897932) and *DDX39B* (rs2523506) that enhances sIL7R expression, leading to approximately three times higher MS risk (Galarza-Munoz et al., 2017). Studies of Mackall and colleagues provided evidence of causality of sIL7R in MS and a potential mechanism of action (Lundstrom et al., 2013). These authors showed that co-administration of recombinant sIL7R and IL7 in mice exacerbated the severity, progression and overall disability of the MS-like disease in the Experimental Autoimmune Encephalomyelitis (EAE) model of MS (Lundstrom et al., 2013). Mechanistically, they showed that sIL7R potentiates the bioavailability and activity of IL7 *in vivo*, leading to enhanced peripheral expansion of CD4^+^ and CD8^+^ T cells (Lundstrom et al., 2013). Such potentiation of T cell survival and proliferation has also been shown by *in vitro* treatment of CD8^+^ T cells with sIL7R/IL7 complexes (Cote et al., 2015). The IL7/IL7R signaling pathway has been shown to preferentially promote expansion of myelin-specific T cells (Bebo et al., 2000; Bielekova et al., 1999; Traggiai et al., 2001), and inhibition of this signaling pathway suppresses the encephalitogenicity of myelin-specific CD4^+^ T cells (Nuro-Gyina et al., 2016). Therefore, by potentiating IL7, it is likely that sIL7R preferentially expands the population of T cells that destroy myelin in MS. In addition to MS, sIL7R has been reported to be elevated in patients of other autoimmune disorders, including type 1 diabetes, rheumatoid arthritis and systemic lupus erythematosus, and to positively correlate with disease activity (Badot et al., 2011; Badot et al., 2013; Lauwerys et al., 2014; Monti et al., 2013). Collectively, this body of work supports a role of sIL7R in the development of MS and other autoimmune diseases.

Given that alternative splicing of exon 6 is the main determinant of sIL7R expression in humans, we identified this process as a potential therapeutic target, and one that could be dialed down (decrease exon 6 exclusion and sIL7R) to reduce autoimmunity, or dialed up (increase exon 6 exclusion and sIL7R) if self-reactivity is desired, as in immuno-oncology. This tuning of sIL7R expression could be achieved using splicing-modulating (also known as splice-switch) antisense oligonucleotides (ASOs). ASOs hold great promise by enabling therapeutic targeting of previously “undruggable” disease-causing genes but this promise was hindered for decades by the poor pharmacodynamics of these nucleic acid-based drugs. Recent developments in oligonucleotide chemistry and delivery technologies have improved the pharmacological properties of ASOs and empowered their clinical potential (Kulkarni et al., 2021; Roberts et al., 2020; Seth et al., 2019). These efforts have led to the approval of several splicing-modulating ASOs by the FDA, including eteplirsen (Exondys 51^®^), golodirsen (Vyondys 53^®^) and casimersen (Amondys 45^®^) for treatment of different subsets of Duchenne muscular dystrophy patients (Charleston et al., 2018; Cirak et al., 2011; Frank et al., 2020; Mendell et al., 2016; Wagner et al., 2021), and nusinersen (Spinraza^®^) for all types of spinal muscular atrophy (SMA). Nusinersen best exemplifies the therapeutic potential of splicing-modulating ASOs. SMA is caused by insufficient production of the survival motor neuron (SMN) protein due to: (*i*) disruptive mutations in the SMN1 gene (Lunn and Wang, 2008), and (*ii*) a polymorphism in the almost identical SMN2 gene that promotes exclusion of the seventh exon from SMN2 transcripts, leading to production of a truncated, inactive protein (Kolb and Kissel, 2011). Nusinersen rescues production of functional, full-length SMN protein by promoting inclusion of exon 7 in SMN2 pre-mRNAs, leading to significant improvement or stabilization of otherwise progressively deteriorating motor function in SMA patients (Darras et al., 2019; De Vivo et al., 2019; Finkel et al., 2016; Finkel et al., 2017; Mercuri et al., 2018; Pera et al., 2021). The clinical success of Nusinersen, together with the pro-immune effects of sIL7R, prompted us to develop ASOs that control sIL7R levels by modulating IL7R exon 6 splicing.

We have extensively studied the splicing of IL7R exon 6, which led to the characterization of numerous *cis*-acting splicing elements and *trans*-acting splicing factors that determine the inclusion or exclusion of the exon, and consequently set the levels of sIL7R (Evsyukova et al., 2013; Galarza-Munoz et al., 2017; Gregory et al., 2007; Schott et al., 2021). We uncovered two exonic splicing enhancers (ESEs) within exon 6 that promote exon inclusion, and three exonic splicing silencers (ESSs) that promote exon exclusion (Evsyukova et al., 2013), among them the enhancer ESE2 (nt 36-40 of exon 6) and the silencer ESS3 (nt 51-65 of exon 6), which are relevant to work conducted here. Among *trans*-acting splicing factors, we uncovered the RNA helicase DDX39B as an activator of the exon, which requires an intact ESE2 to promote its inclusion (Galarza-Munoz et al., 2017). We also identified several repressors of the exon including CPSF1 (Evsyukova et al., 2013), PTBP1 and U2AF2 (Schott et al., 2021), the latter of which explains the mechanism by which the SNP rs6897932 influences exon 6 splicing. This insight into the network of elements/factors regulating splicing of IL7R exon 6 facilitated the rational design of ASOs to control its splicing.

In this study we uncovered two anti-sIL7R ASOs and two pro-sIL7R ASOs, and showed that these ASOs effectively modulate splicing of exon 6 and sIL7R secretion in cell lines and in human primary CD4^+^ T cells. Underscoring their therapeutic potential, we showed these ASOs correct (anti-sIL7R ASOs) or phenocopy (pro-sIL7R ASOs) the effects of the MS risk allele of rs6897932 on exon 6 splicing. This proof-of-concept study established the effectiveness of these ASOs to control IL7R splicing and sIL7R secretion in human T cells and future pre-clinical studies will assess their potential for therapeutic intervention in autoimmunity and immuno-oncology.

## RESULTS

### Targeted ASO screen identified functional IL7R ASOs

To facilitate discovery of antisense oligonucleotides (ASOs) that promote inclusion of IL7R exon 6, we created a GFP-based fluorescent reporter of IL7R exon 6 splicing. This GFP-IL7R fluorescent reporter was generated by cloning the genomic sequence of IL7R spanning the distal portion of intron 5 (614 nt), exon 6 (94 nt) and the proximal portion of intron 6 (573 nt) interrupting the coding sequence of GFP. In this reporter, GFP expression is controlled by splicing of IL7R exon 6: exclusion of exon 6 splices together the coding sequence of GFP leading to GFP expression, whereas exon 6 inclusion causes a shift in the reading frame that abolishes GFP expression (Fig. 1A). Accordingly, GFP expression from the reporter is produced directly proportional to the levels of exon 6 exclusion. We created a HeLa cell line stably expressing an inducible version of the GFP-IL7R reporter and used this stable cell line to identify ASOs that promote inclusion or exclusion of exon 6 via quantification of GFP expression by flow cytometry.

**Figure 1.**
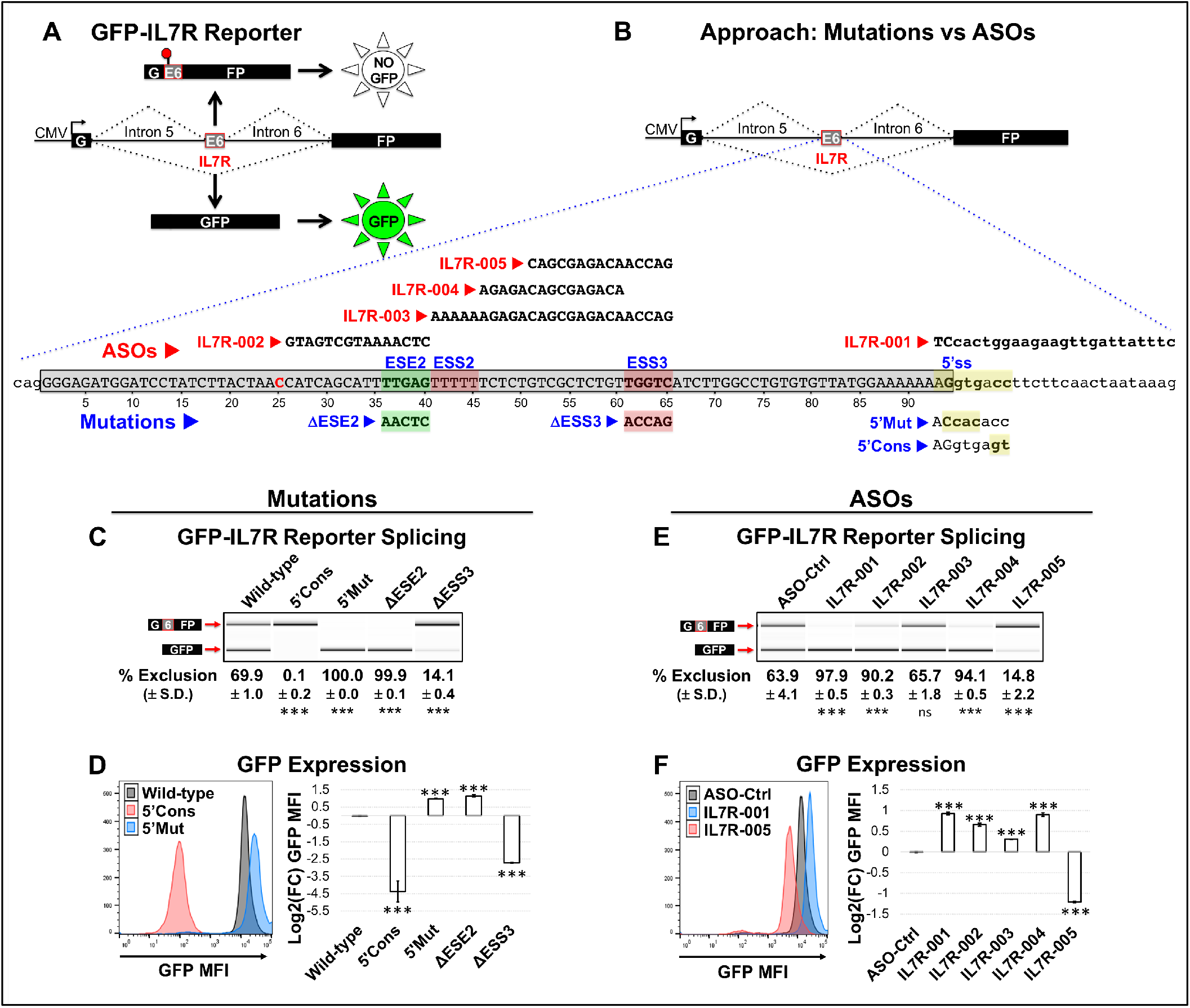
Screen of ASOs targeting functional *cis*-acting splicing elements within IL7R exon 6. *(A)* Schematics of the GFP-IL7R splicing reporter, wherein GFP is only expressed by exclusion of IL7R exon 6. *(B)* Schematics of approach comparing the effects of mutagenesis of functional *cis*-acting splicing elements to steric blocking of the corresponding elements with ASOs using the GFP-IL7R reporter. The blue dotted lines expand the exon 6 sequence (gray box), and the relevant *cis*-acting elements are color-coded: enhancer ESE2 (green), silencers ESS2 and ESS3 (red), and 5’ss (yellow). Mutations to ESE2, ESS3 and the 5’SS are shown underneath each element: *Δ*ESE2 and *Δ*ESS3 represent mutations to ESE2 and ESS3, respectively, and 5’Cons and 5’Mut represent consensus and crippling mutations to the 5’ss, respectively. The sequence of ASOs targeting these *cis*-elements (IL7R-001 – IL7R-005) is shown above the exon sequence: IL7R-001 blocks the 5’ss of exon 6, IL7R-002 blocks ESE2, IL7R-003 blocks ESS2 and ESS3, IL7R-004 blocks the sequence in between ESS2 and ESS3, and IL7R-005 blocks ESS3. *(C, E)* Percentage (%) of exon 6 exclusion (mean ± S.D.) in transcripts from the reporter in HeLa cells stably expressing wild-type or mutant versions of the reporter (*C*), or cells stably expressing the wild-type reporter and transfected with control (ASO-Ctrl) or experimental (IL7R-001 – IL7R-005) morpholino ASOs (*E*) are shown under the gel images. The top band in the gels represents the exon 6 included product whereas the lower band represents the exon 6 excluded product. *(D, F)* Mean fluorescence intensity (MFI) of GFP expression in the cells from panel *C* stably expressing the different versions of the reporter (*D*), or in cells from panel *E* transfected with ASOs (*F*). The left panels show representative GFP MFI histograms for selected mutants (Wild-type, 5’Cons and 5’Mut) (*D*) and selected ASOs (ASO-Ctrl, IL7R-001 and IL7R-005) (*F*), with quantification shown on the right as Log2(Fold-Change) [Log2(FC)] relative to wild-type for all mutants (*D*) and relative to control for all ASOs (*F*). (***) *p < 0.001*.

To test the feasibility of the ASO approach to control splicing of IL7R exon 6, we compared the effects of ASO-mediated steric blocking of specific *cis*-acting splicing elements within exon 6 to the effects of mutations in the corresponding *cis*-acting elements (Fig. 1B). We first focused on the weak 5’-splice site (5’ss) of exon 6 (AGgtga<u>cc</u>). We introduced mutations to the 5’ss in the reporter to either make it a consensus 5’ss (5’Cons; AGgtga<u>gt</u>) or to cripple it (5’Mut; A<u>Ccac</u>acc) (Fig. 1B), and assessed the effects of these mutations on exon 6 splicing and GFP expression. All versions of the reporter contain the risk ‘C’ allele at the SNP rs6897932. While exon 6 was excluded in 70% of the transcripts from the wild-type reporter, the consensus mutation (5’Cons) caused complete inclusion of the exon, and the crippling mutation (5’Mut) caused complete exclusion of the exon (Fig. 1C). As expected, the 5’Cons mutation abolished GFP expression, whereas the 5’Mut led to a 2-fold increase in GFP expression [equivalent to a Log2(fold-change) of 0.99; hereafter referred to as Log2(FC)] (Fig. 1D). These results showed that GFP expression is a reliable proxy of exon 6 splicing in the reporter and established the range of GFP expression from the reporter.

We next tested whether a morpholino ASO blocking the 5’ss (IL7R-001) could phenocopy the effect of the crippling 5’ss mutation (5’Mut) (Fig. 1B). To this end, we transfected 10 µM of IL7R-001 or a non-targeting control morpholino ASO (ASO-Ctrl) into the wild-type reporter cell line using the Endo-Porter transfection reagent (Gene Tools) and assayed cells for IL7R splicing and GFP expression as before. Similar to the crippling 5’Mut mutation, ASO IL7R-001 caused complete exon 6 exclusion (Fig. 1E), leading to a 1.9-fold increase in GFP expression [Log2(FC) = 0.93] (Fig. 1F). This analysis showed that blocking of the 5’ss with ASO IL7R-001 reproduced the effect of the crippling 5’ss mutation, thereby validating the effectiveness of ASOs to control exon 6 splicing.

In addition to the 5’ss, we also targeted the previously identified ESE2 (nt 36-40 of exon 6; Fig. 1B) (Evsyukova et al., 2013; Galarza-Munoz et al., 2017). As expected, mutation of ESE2 (*Δ*ESE2) in the GFP-IL7R reporter entirely excluded the exon (Fig. 1C) and enhanced GFP expression by 2.2-fold [Log2(FC) = 1.1] (Fig. 1D) compared to the wild-type reporter. These effects were largely reproduced by a morpholino ASO targeting ESE2 (IL7R-002, complementary to nt 26-40; Fig. 1B), which enhanced exon 6 exclusion to 90.2% (Fig. 1E) and increased GFP expression by 1.6-fold [Log2(FC) = 0.66] (Fig. 1F) compared to the control ASO (ASO-Ctrl). Collectively, the results of IL7R-001 and IL7R-002 showed that splicing of IL7R exon 6 can be effectively modulated with ASOs targeting either core splicing elements like the 5’ss (IL7R-001) or auxiliary splicing elements like ESE2 (IL7R-002).

Having established the utility of ASOs to control splicing of IL7R exon 6, we focused on our primary goal of identifying ASOs that reduce exon 6 exclusion. Systematic mutagenesis of exon 6 previously identified several silencing elements within nt 41-65, among them ESS2 between nt 41-45 and ESS3 between nt 51-65 (Evsyukova et al., 2013). We designed the ASO IL7R-003 targeting nt 41-65, which blocks both ESS2 and ESS3 (Fig. 1B). Surprisingly, transfection of IL7R-003 into the wild-type reporter cell line had negligible effects on exon 6 splicing (Fig. 1E) and GFP expression (Fig. 1F) compared to the control ASO. This suggests the presence of a previously unknown enhancer within this target sequence, most likely between nt 46-50. To further dissect this, we created ASOs IL7R-004 targeting nt 46-60, and IL7R-005 targeting nt 51-65 (Fig. 1B). While these two ASOs target a shared core sequence between nt 50-60, IL7R-004 additionally targets a potential enhancer in nt 46-50, and IL7R-005 additionally targets a predicted silencer in nt 61-65. IL7R-004 was found to promote almost complete exon 6 exclusion (Fig. 1E) leading to high levels of GFP expression (1.9-fold increase; [Log2(FC) = 0.90]) (Fig. 1F), thereby confirming the likely presence of an enhancer between nt 46-50. On the other hand, IL7R-005 was found to largely reduce exon 6 exclusion (Fig. 1E) leading to low levels of GFP expression (2.3-fold reduction; [Log2(FC) = −1.20]) (Fig. 1F). Indeed, the effects of IL7R-005 were comparable to mutation of nt 61-65 of ESS3 (*Δ*ESS3), which almost abolished exon 6 exclusion (Fig. 1C) and GFP expression (6.5-fold reduction; [Log2(FC) = −2.71]) (Fig. 1D). In addition to these splicing elements, we have also identified an intronic splicing silencer (ISS1) consisting of a polyadenylation signal (pA; AAUAAA) in the vicinity of the 5’ss in intron 6, which silences splicing of exon 6 most likely because binding of CPSF1 to this pA signal impedes U1 snRNP binding to the adjacent 5’ss (Evsyukova et al., 2013). An ASO that blocks ISS1, IL7R-006, also reduced exon 6 exclusion in the reporter (Fig. S1A) and GFP expression (Fig. S1B).

The results above showed that ASOs blocking critical *cis*-acting splicing elements in exon 6 or intron 6 largely phenocopy the effects of mutation of the corresponding elements, thereby validating the use of ASOs to control IL7R splicing. Most importantly, this targeted screen uncovered ASOs IL7R-005 and IL7R-006 that reduce exon 6 exclusion, and ASOs IL7R-001 and IL7R-004 that enhance exon 6 exclusion.

### ASOs modulate IL7R splicing in a concentration-dependent fashion

After discovering ASOs that modulate exon 6 splicing we interrogated whether or not these ASOs could do so in a concentration-dependent manner. To evaluate the response of IL7R exon 6 splicing to ASO concentration, control (ASO-Ctrl) or experimental (IL7R-001, IL7R-004, IL7R-005 and IL7R-006) morpholino ASOs were transfected into the wild-type reporter cell line at different concentrations (0, 1, 5, and 10 µM), and the cells were assayed for GFP expression by flow cytometry 48 hours after. Representative histograms of GFP expression are shown for each ASO at the various concentrations in the left panel of Figure 2A. GFP expression was quantified as mean fluorescence intensity (MFI) and converted to Log2(FC) relative to 0 µM and are shown as a function of ASO concentration in the right panel of Figure 2A. While increasing concentrations of ASO-Ctrl did not alter GFP expression, increasing concentrations of the experimental IL7R ASOs induced significant changes in GFP expression. Increasing concentrations of ASOs IL7R-001 and IL7R-004 progressively increased GFP expression reaching 0.81 and 0.84 Log2(FC), respectively, at the highest dose tested (Fig. 2A, right panel). On the other hand, increasing concentrations of ASOs IL7R-005 and IL7R-006 progressively reduced GFP expression to −1.08 and −0.71 Log2(FC), respectively, at the highest dose (Fig. 2A, right panel). These changes in GFP expression imply a concentration-dependent modulation of exon 6 splicing in the reporter. Indeed, RT-PCR analysis of exon 6 splicing in transcripts from the reporter confirmed a lack of splicing modulation by increasing concentrations of ASO-Ctrl, a gradual increase in exon 6 exclusion by ASOs IL7R-001 and IL7R-004, and a gradual decrease in exon 6 exclusion by ASOs IL7R-005 and IL7R-006 (Fig. 2B). All experimental IL7R ASOs showed concentration-dependent modulation of GFP expression and exon 6 splicing, with IL7R-001 and IL7R-004 increasing exon 6 exclusion with similar potency, whereas IL7R-005 decreased exon 6 exclusion with slightly more potency than IL7R-006.

**Figure 2.**
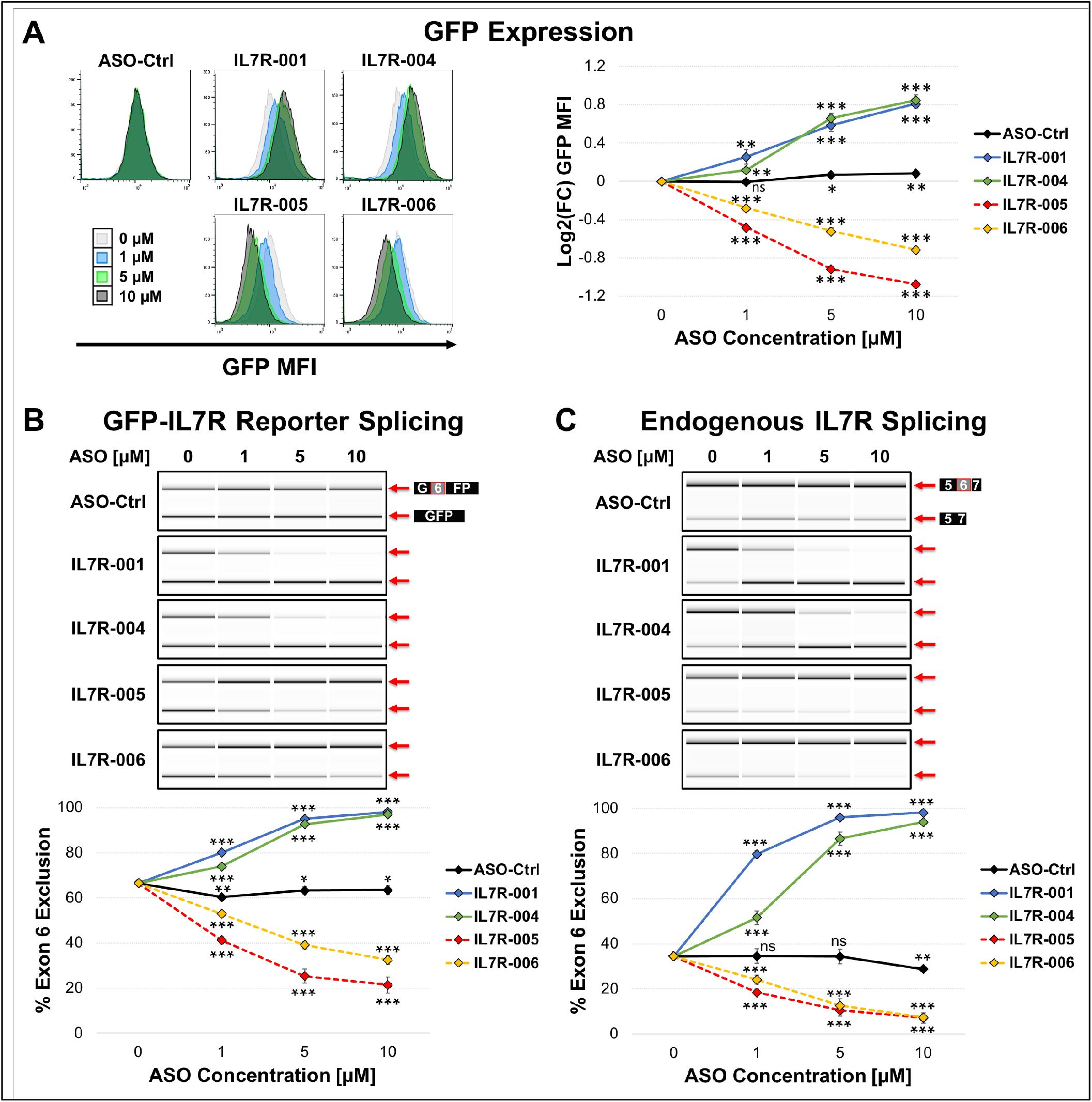
Concentration-dependent effect of lead ASOs on IL7R exon 6 splicing. *(A)* GFP expression in the wild-type reporter cell line transfected with increasing concentrations of control (ASO-Ctrl) or experimental (IL7R-001, IL7R-004, IL7R-005 and IL7R-006) morpholino ASOs. Overlapped representative histograms are shown on the left for each ASO at 0 μM (light gray), 1 μM (blue), 5 μM (green) and 10 μM (dark gray), with quantification of MFI plotted on the right as dose-response curves of Log2(FC) relative to 0 µM for ASO-Ctrl (black), IL7R-001 (blue), IL7R-004 (green), IL7R-005 (red) and IL7R-006 (yellow). *(B-C)* Percentage (%) of exon 6 exclusion in transcripts from the GFP-IL7R reporter (*B*) or the endogenous IL7R gene (*C*) Representative gel images for each ASO are shown at the top with quantification of percent exon 6 exclusion (mean ± S.D.) plotted at the bottom as a function of ASO concentration (color-coded as in *A*). (*) *p < 0.05*, (**) *p < 0.01*, and (***) *p < 0.001*.

The translatability of these ASOs for therapeutic applications requires modulation of exon 6 splicing in transcripts from the endogenous *IL7R* gene. Although *IL7R* is predominantly expressed in lymphocytes, its expression is up-regulated in many cancer cell lines, including HeLa cells, and we have previously shown that IL7R exon 6 is similarly regulated in HeLa cells, Jurkat T cells and human primary CD4^+^ T cells (Galarza-Munoz et al., 2017). Accordingly, we next examined the concentration-dependent effect of the ASOs on exon 6 splicing of the endogenous IL7R transcripts. In HeLa cells, exon 6 is excluded in 34.4% of the endogenous IL7R transcripts, and this percentage remained constant with increasing concentrations of ASO-Ctrl (Fig. 2C). Conversely, increasing concentrations of IL7R-001 and IL7R-004 progressively enhanced exon 6 exclusion in the endogenous IL7R transcripts to 98.1 and 93.9 %, respectively (Fig. 2C). Although these ASOs had similar effects at the highest concentration, IL7R-001 was more effective at lower concentrations. Conversely, increasing concentrations of IL7R-005 and IL7R-006 gradually reduced exon 6 exclusion in the endogenous IL7R transcripts, with both ASOs being equally potent in promoting exon 6 inclusion in the endogenous IL7R transcripts (7.1 and 7.4 %, respectively, at the highest concentration) (Fig. 2C). Collectively, these analyses showed that our lead IL7R ASOs modulate exon 6 splicing in transcripts from both the reporter and the endogenous IL7R gene, and that the effect on exon 6 splicing can be dialed up or down in a concentration-dependent manner.

### ASOs modulate secretion of sIL7R

We next assessed the effect of these ASOs on the secretion of sIL7R. IL7R-001 and IL7R-004 increase exclusion of exon 6 and thus are predicted to enhance sIL7R secretion, whereas ASOs IL7R-005 and IL7R-006 decrease exclusion of exon 6 and are predicted to reduce sIL7R secretion. We first tested these predictions in HeLa cells, which regulate exon 6 splicing similarly than human T cells (Galarza-Munoz et al., 2017). Control (ASO-Ctrl) or experimental (IL7R-001, IL7R-004, IL7R-005 and IL7R-006) morpholino ASOs were transfected into HeLa cells at 10 µM with the Endo-Porter transfection reagent, and cells were assayed three days after for exon 6 splicing and sIL7R secretion. RT-PCR analysis of transfected cells revealed 31.7% exon 6 exclusion in cells transfected with ASO-Ctrl, and this was largely increased by ASOs IL7R-001 (96.0%) and IL7R-004 (86.1%), and decreased by ASOs IL7R-005 (13.9%) and IL7R-006 (17.7%) (Fig. S2A). As predicted, secretion of sIL7R was modulated by the ASOs proportionally to their effect on exon 6 exclusion. ASOs IL7R-001 and IL7R-004 increased sIL7R secretion by 1.8-fold and 1.9-fold, respectively, and ASOs IL7R-005 and IL7R-006 decreased sIL7R secretion by 2.2-fold and 2.4-fold, respectively (Fig. S2B). These results showed that lead morpholino IL7R ASOs modulate the expression of sIL7R when transfected into HeLa cells.

IL7R is predominantly expressed in T cells, and thus these are the major producers of sIL7R *in vivo*. Accordingly, we next evaluated the effect of lead ASOs on the secretion of sIL7R in human primary CD4^+^ T cells isolated from healthy donors. The ASOs were transfected into the primary T cells by nucleofection (Distler et al., 2005) and the cells were assayed three days after for exon 6 splicing and sIL7R secretion. While exon 6 was excluded in 17.0% of IL7R transcripts in T cells transfected with ASO-Ctrl, it was completely excluded with ASOs IL7R-001 and IL7R-004, and nearly completely included with ASOs IL7R-005 and IL7R-006 (Fig. 3A). This modulation of exon 6 splicing led to the expected changes in the secretion of sIL7R. ASOs IL7R-001 and IL7R-004 significantly increased sIL7R secretion (4.5-fold and 3.8-fold, respectively), whereas ASOs IL7R-005 and IL7R-006 decreased sIL7R secretion by greater than 5-fold (reduced to near or below the limit of detection) (Fig. 3B). Since ASOs IL7R-001 and IL7R-004 enhance sIL7R secretion, these are hereafter collectively referred to as pro-sIL7R ASOs. ASOs IL7R-005 and IL7R-006 reduce sIL7R secretion, and thus are hereafter collectively referred to as anti-sIL7R ASOs. Together, these results showed that lead IL7R ASOs modulate splicing of exon 6 in human CD4^+^ T cells and enable control of sIL7R expression in this biologically relevant cell type. Given the proposed role of sIL7R in the expansion of T cells and the development of autoimmunity (Cote et al., 2015; Lundstrom et al., 2013), these IL7R ASOs have potential therapeutic utilities in autoimmunity and cancer.

**Figure 3.**
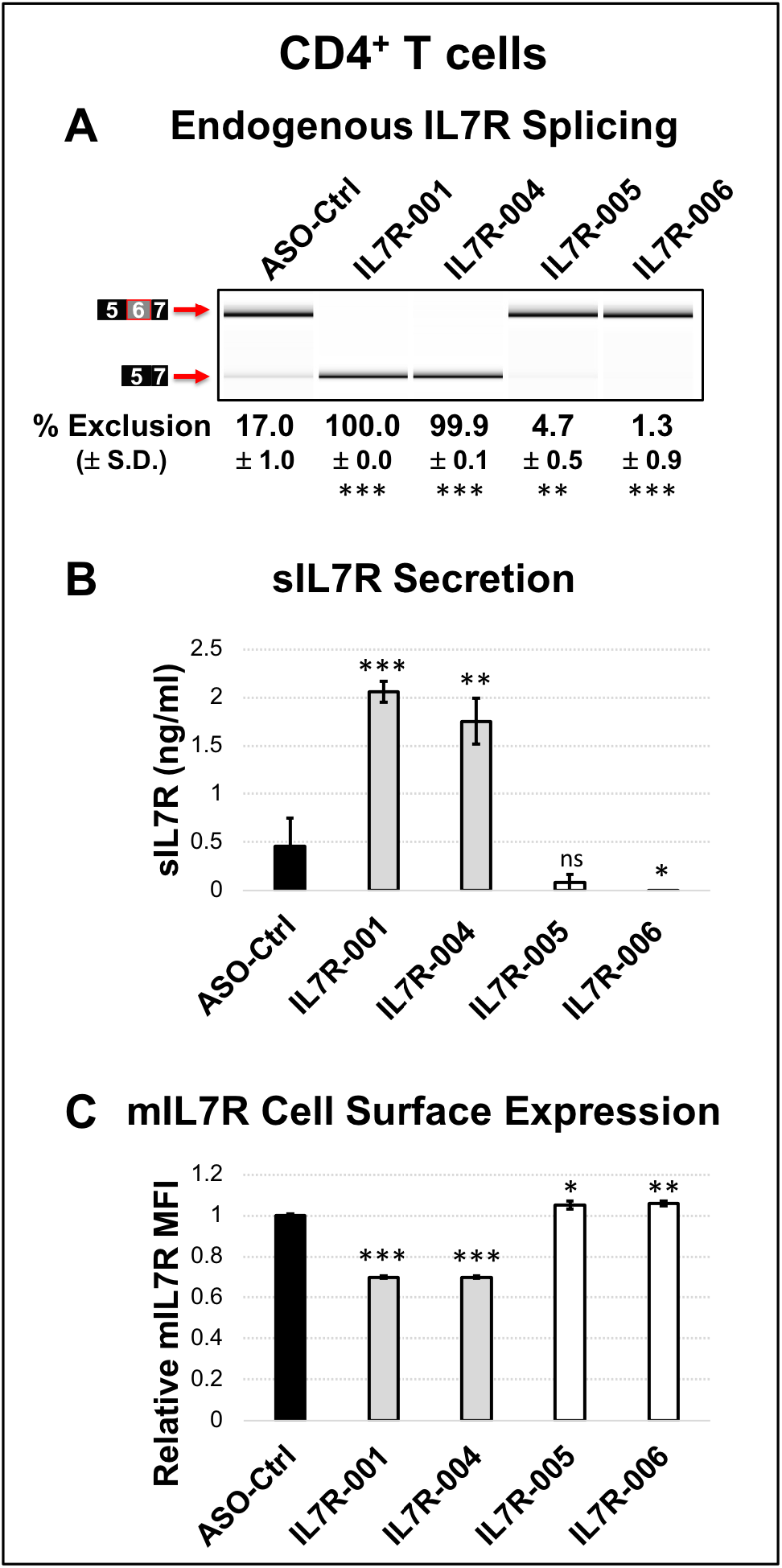
Lead ASOs modulate expression of IL7R protein isoforms in human primary CD4^+^ T cells. *(A)* Representative gel image of exon 6 splicing in the endogenous IL7R transcripts from human primary CD4^+^ T cells transfected with control (ASO-Ctrl) or experimental (IL7R-001, IL7R-004, IL7R-005 and IL7R-006) morpholino ASOs. Percent (%) exon 6 exclusion (mean ± S.D.) is shown for each ASO under the gel image. *(B)* Levels of secreted sIL7R (mean ± S.D.) in supernatants from cells in panel *A*. *(C)* Relative MFI values (mean ± S.D.) of mIL7R cell surface expression in cells from panel *A*. (*) *p < 0.05*, (**) *p < 0.01*, and (***) *p < 0.001*.

We next examined the cell surface expression of mIL7R given that exon 6 exclusion increases sIL7R RNA expression at the expense of mIL7R RNA. This is important because loss of mIL7R function in mice and humans causes lymphopenia and severe immunodeficiency (Maraskovsky et al., 1996; Peschon et al., 1994; Puel et al., 1998; Roifman et al., 2000), and thus therapies that inhibit the expression and/or function of mIL7R could cause broad immunosuppression. As expected, anti-sIL7R ASOs IL7R-005 and IL7R-006 slightly increased mIL7R cell surface expression in primary CD4^+^ T cells (Fig. 3C), and thus are predicted to avoid immunosuppression. Cell surface expression of mIL7R in the primary CD4^+^ T cells treated with pro-sIL7R ASOs IL7R-001 and IL7R-004 was reduced ∼30% compared to T cells treated with control ASO (Fig. 3C). This reduction is not profound and is predicted to be tolerable since heterozygous knockout of IL7R in mice is tolerated (Peschon et al., 1994). These results showed that IL7R ASOs provide excellent control of sIL7R levels with minor effects on the cell surface expression of mIL7R.

### Anti-sIL7R ASOs correct increased exclusion of exon 6 caused by the MS risk allele of the SNP rs6897932

Human genetic and molecular studies have revealed the MS risk SNP rs6897932 as the main determinant of sIL7R expression, with the risk ‘C’ allele enhancing exon 6 exclusion and sIL7R secretion (Gregory et al., 2007; Hoe et al., 2010; Lundstrom et al., 2013). Therefore, in order for anti-sIL7R ASOs to be therapeutic in MS, they will need to reduce the enhanced exclusion of exon 6 caused by the risk ‘C’ allele of rs6897932 to levels equal or smaller than those of the protective “T” allele. To assess this, we used versions of the GFP-IL7R reporter carrying either the protective ‘T’ allele (T reporter) or the risk ‘C’ allele (C reporter) of rs6897932. We transfected ASO-Ctrl, IL7R-005 or IL7R-006 morpholino ASOs into HeLa cells stably expressing the C reporter, and assessed the effects of the ASOs on exon 6 splicing compared to cells stably expressing the T reporter and transfected with ASO-Ctrl. Illustrating the effects of the SNP, RT-PCR analysis in cells treated with ASO-Ctrl showed that exon 6 was excluded at a significantly higher frequency (78.2% exclusion) in the C reporter cell line than in the T reporter cell line (57.2% exclusion) (Fig. 4A). Importantly, anti-sIL7R ASOs IL7R-005 and IL7R-006 restored exon 6 exclusion in the C reporter cell line to levels similar or lower than those observed in the T reporter cell line, with IL7R-005 (44.3% exclusion) showing more potency than IL7R-006 (62.8% exclusion) (Fig. 4A). Because GFP expression from the reporter is a proxy for sIL7R, we also examined effects of the ASOs on GFP expression. As expected, the anti-sIL7R ASOs reduced GFP expression to levels equal or lower than those in the T reporter cell line (Fig. 4B), implying that anti-sIL7R ASOs would also correct sIL7R expression. These results showed that anti-sIL7R ASOs IL7R-005 and IL7R-006 effectively correct the enhanced exclusion of IL7R exon 6 caused by the risk ‘C’ allele of rs6897932, thereby demonstrating their utility to control splicing of this critical exon for potential treatment of MS patients with high levels of sIL7R.

**Figure 4.**
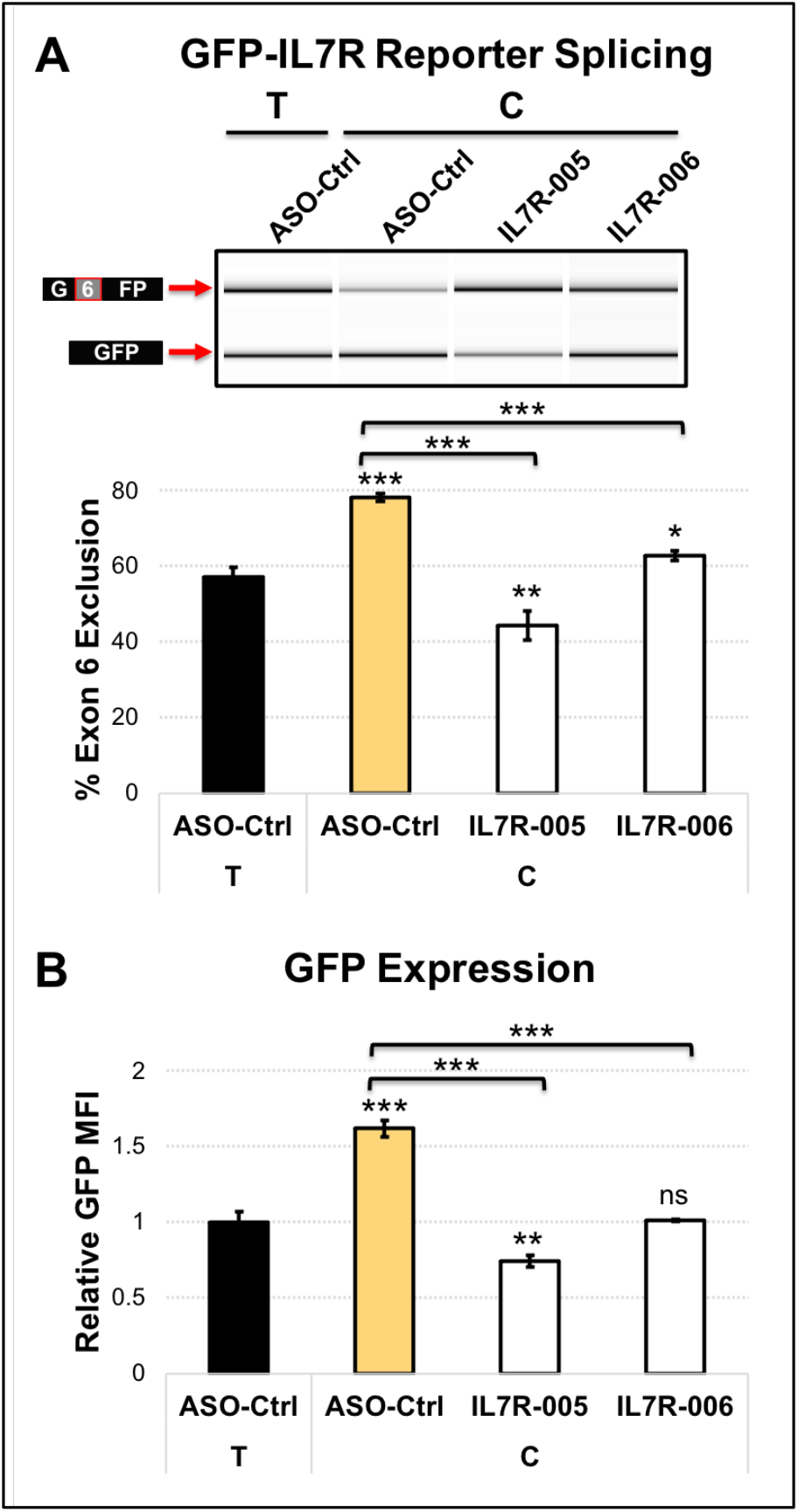
Anti-sIL7R ASOs correct the enhanced exclusion of IL7R exon 6 in the presence of the MS risk ‘C’ allele of rs6897932. Control (ASO-Ctrl) or lead anti-sIL7R morpholino ASOs IL7R-005 and IL7R-006 were transfected at 10 µM with Endo-Porter into HeLa cells stably expressing versions of the GFP-IL7R reporter containing either the protective ‘T’ allele or the risk ‘C’ allele of the MS SNP rs6897932. *(A)* Representative gel image of exon 6 splicing in transcripts from the GFP-IL7R reporter containing either the protective ‘T’ allele or the risk ‘C’ allele of the MS SNP rs6897932, and transfected with control (ASO-Ctrl) or lead anti-sIL7R (IL7R-005 and IL7R-006) morpholino ASOs. Percentage of exon 6 exclusion (mean ± S.D.) for each condition is plotted below the gel image. *(B)* GFP MFI (mean ± S.D.) from cells in panel *A* shown relative to the T reporter treated with ASO-Ctrl. (*) *p < 0.05*, (**) *p < 0.01*, and (***) *p < 0.001*.

### Pro-sIL7R ASOs phenocopy increased exclusion of exon 6 caused by the MS risk allele of the SNP rs6897932

Considering the role of sIL7R in enhancing self-reactive immune responses, and the fact that immune responses against cancer cells are self-reactive in nature, we propose that up-regulation of sIL7R could be used to enhance anti-cancer immunity. Given that the SNP rs6897932 is a key determinant of sIL7R expression, we predict that to effectively enhance anti-cancer immunity, sIL7R needs to be raised to or above the levels observed in individuals homozygous for the risk ‘C’ allele at rs6897932. To assess this, we evaluated whether pro-sIL7R ASOs could enhance the lower levels of exon 6 exclusion in the T reporter to or above the levels observed in the C reporter. Control (ASO-Ctrl) and pro-sIL7R ASOs IL7R-001 and IL7R-004 morpholino ASOs were transfected into cells stably expressing the T reporter, and the cells were assayed for IL7R exon 6 exclusion and GFP expression relative to cells stably expressing the C reporter treated with ASO-Ctrl. As expected, higher exon 6 exclusion was observed in the C reporter cells treated with ASO-Ctrl (78.0% exclusion) than in the T reporter cells treated with ASO-Ctrl (54.4% exclusion) (Fig. 5A). Pro-sIL7R ASOs IL7R-001 and IL7R-004 elevated exon 6 exclusion in the T reporter cells to 91.5% and 88.3%, respectively, both of which are significantly higher than the 78.0% observed in the C reporter cells treated with ASO-Ctrl (Fig. 5A). GFP expression in the T reporter cells treated with IL7R-001 and IL7R-004 was significantly higher (∼1.4 fold) than in the C reporter cells treated with ASO-Ctrl (Fig. 5B), thereby implying that these ASOs can surpass the targeted level of sIL7R expression imposed by the risk ‘C’ allele. These results showed that pro-sIL7R ASOs IL7R-001 and IL7R-004 can enhance exclusion of IL7R exon 6 to levels above those observed in individuals with high predisposition to self-reactivity, and thus are good candidates to test the premise that up-regulation of sIL7R can boost anti-cancer immunity.

**Figure 5.**
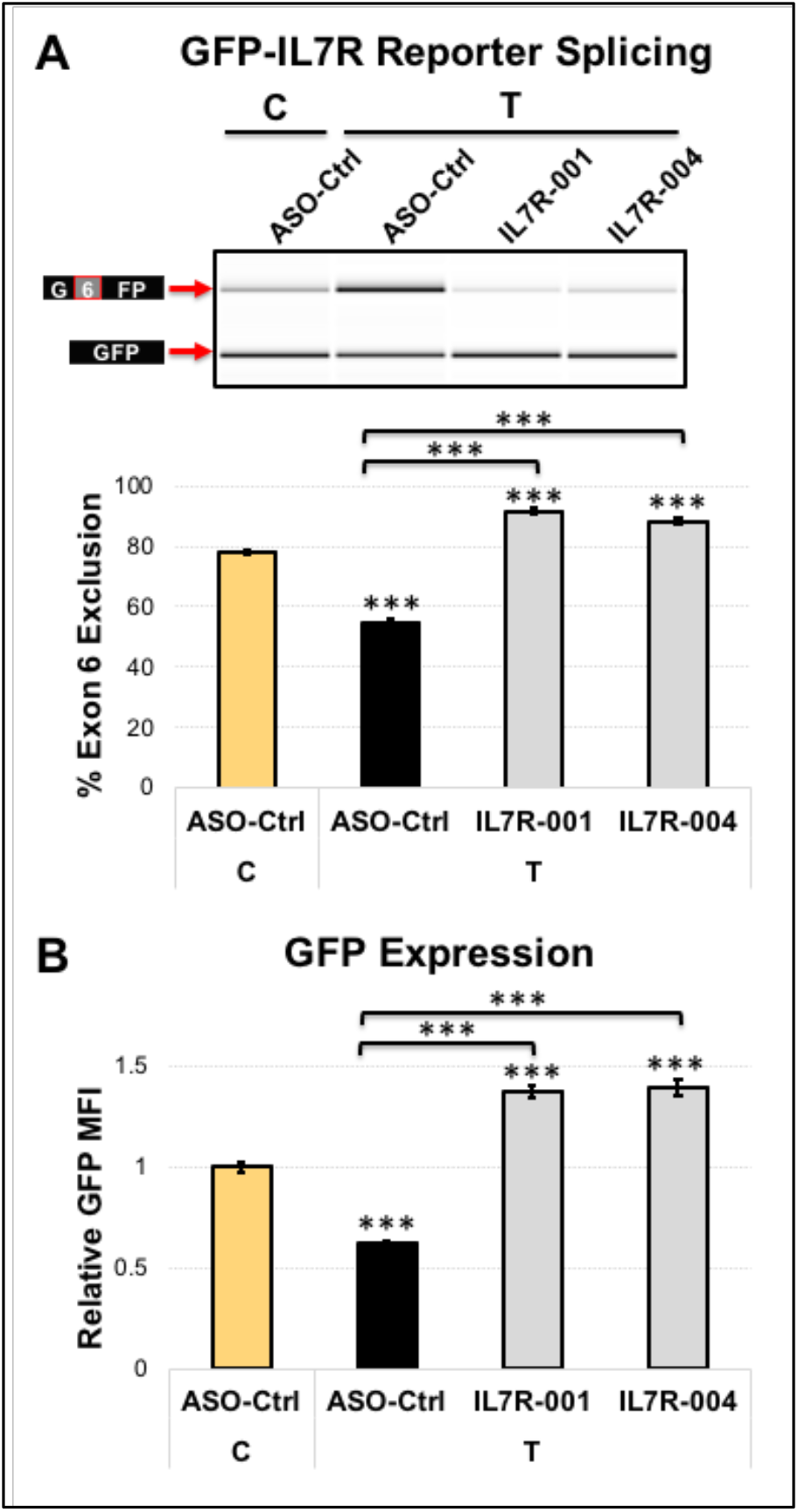
Pro-sIL7R ASOs enhance exon 6 exclusion in the presence of the protective ‘T’ allele of the SNP rs6897932. *(A)* Representative gel image of exon 6 splicing in transcripts from the GFP-IL7R reporter containing either the MS risk ‘C’ allele or the protective ‘T’ allele of the IL7R SNP rs6897932, and transfected with control (ASO-Ctrl) or lead pro-sIL7R (IL7R-001 and IL7R-004) morpholino ASOs. Percentage of exon 6 exclusion (mean ± S.D.) for each condition is plotted below the gel image. *(B)* GFP MFI (mean ± S.D.) from cells in panel *A* shown relative to the C reporter treated with ASO-Ctrl. (***) *p < 0.001*.

## DISCUSSION

Elevated expression of sIL7R is a driver of MS and autoimmunity. Since the expression of sIL7R is determined by exclusion of exon 6, the goal of this study was to identify ASOs that control the expression of sIL7R by modulating splicing of this exon that could be ultimately developed as novel immunotherapeutic drugs. Accomplishing this goal, we discovered the anti-sIL7R ASOs IL7R-005 and IL7R-006, which reduce sIL7R secretion by promoting inclusion of exon 6, and the pro-sIL7R ASOs IL7R-001 and IL7R-004, which enhance sIL7R secretion by promoting exclusion of the exon. The main determinant of sIL7R expression is the risk allele of the MS risk SNP rs6897932 (Gregory et al., 2007; Hoe et al., 2010; Lundstrom et al., 2013), and we further showed that anti-sIL7R ASOs correct the enhanced exclusion of exon 6 caused by the risk allele, whereas pro-sIL7R ASOs phenocopy this effect. Accordingly, these ASOs are excellent candidates to control sIL7R expression for therapeutic purposes in autoimmunity and immuno-oncology.

We postulate the anti-sIL7R ASOs IL7R-005 and IL7R-006 would provide an effective and safe therapy for MS, which is a critical unmet need. Treatment of MS has been considerably challenging due to the multi-factorial origin of the disease, which results from complex interactions between environmental and genetic factors. Environmental factors include infection with Epstein-Barr virus (EBV) (Bjornevik et al., 2022), vitamin D deficiency (Sintzel et al., 2018), smoking (Degelman and Herman, 2017), and others. Among genetic factors, over 500 putative MS genes have been identified (International Multiple Sclerosis Genetics, 2019), many of which, and likely combinations thereof, could tip the delicate balance between self-tolerance and autoimmunity towards the latter. Independent of which genetic and environmental triggers are at fault, ultimately these putative MS genes enhance self-reactivity for CNS myelin antigens leading to demyelination and neurodegeneration. The development of biologics that could target specific immunological processes to limit the capacity of the immune system to mount immune responses against myelin antigens has greatly improved the efficacy of MS drugs. For example, natalizumab (Tysabri^®^) is a monoclonal antibody that blocks *α*4-integrin in the surface of leukocytes and prevents their migration into tissues, including the CNS (Selewski et al., 2010). Ocrelizumab (Ocrevus^®^) is an anti-CD20 monoclonal antibody that lowers immune activity by depleting CD20-expressing cells, which includes mature B cells (CD19^+^CD20^+^) and a subset of T cells (CD3^+^CD20^+^) (Gingele et al., 2018; Jakimovski et al., 2017). These biological drugs are more efficacious in preventing MS relapses than previous generation of MS drugs, but this improved efficacy comes at the cost of higher adverse side effects largely due to enhanced immunosuppression. These adverse side effects have been particularly damaging during the COVID-19 pandemic; for example, ocrelizumab largely reduces the effectiveness of COVID-19 vaccines (Achiron et al., 2021; Tallantyre et al., 2022), and correlates with higher COVID-19 mortality among MS patients (Prosperini et al., 2022). Accordingly, there exists a need for non-immunosuppressive drugs that are safer for the patients, and we propose this could be achieved by tailoring new drugs to correct specific MS etiologies. The anti-sIL7R ASOs described here approximate this goal by correcting the pathological effects of the MS risk SNP rs6897932 in *IL7R*. These ASOs prevent formation of the pathogenic sIL7R by correcting the increased exclusion of exon 6 caused by the risk allele of this SNP. Exon 6 exclusion and the plasma levels of sIL7R are approximately three times higher in homozygous carriers of the risk “C” allele (CC) than in homozygous carriers of the protective “T” allele (TT) (Lundstrom et al., 2013). Utilizing reporters with the alternative C and T alleles of rs6897932, here we showed anti-sIL7R ASOs restore exon 6 exclusion levels in the C reporter to those observed in the T reporter, with concomitant restoration of GFP expression, which is a proxy for sIL7R expression in these reporters. Accordingly, by correcting exon 6 exclusion in human T cells, these anti-sIL7R ASOs are predicted to restore the plasma levels of sIL7R in CC carriers to those observed in TT carriers. Since these ASOs correct this specific etiology of MS rather than inhibiting immunological processes required for protective immunity against infections and cancers, they are predicted to avoid the adverse immunosuppressive effects observed with other MS therapies.

It is likely that anti-sIL7R ASOs could also be therapeutic in other autoimmune diseases, including type 1 diabetes (T1D), rheumatoid arthritis (RA) and systemic lupus erythematosus (SLE), given sIL7R is elevated in patients from these diseases and correlates with inflammation and/or disease activity (Badot et al., 2011; Badot et al., 2013; Lauwerys et al., 2014; Monti et al., 2013). In support of this, antagonistic anti-IL7R antibodies have been shown to ameliorate and even revert autoimmunity in animal models of MS (Lawson et al., 2015), T1D (Lee et al., 2012; Penaranda et al., 2012), RA (Hartgring et al., 2010), lupus (Gonzalez-Quintial et al., 2011), and inflammatory bowel diseases (Willis et al., 2012). Because these antibodies target the extracellular domain (ECD) of IL7R, and sIL7R is composed of the ECD with a 26 amino acid C-terminus tail, it is likely that these antibodies inhibit sIL7R as well and this contributes to the beneficial effects of these antibodies. Nonetheless, the potential of anti-sIL7R ASOs to be used for treatment of these autoimmune diseases requires further experimentation.

In addition to the anti-sIL7R ASOs described above, we also discovered pro-sIL7R ASOs IL7R-001 and IL7R-004, which increase exon 6 exclusion and sIL7R secretion, and thus are predicted to enhance self-reactivity. While self-reactive immune responses are detrimental in an autoimmune setting, they are instrumental in preventing cancer development and progression. This is the basis of immunotherapies such as immune checkpoint inhibitors, which aim to unleash the capacity of the immune system to eradicate cancers (Pico de Coana et al., 2015; Postow et al., 2015; Sharma and Allison, 2015; Sharma et al., 2021). Although immune checkpoint inhibitors have yielded promising results with some patients reaching full remission, their full potential has been limited by low response rates with only a small fraction of patients exhibiting a response (Haslam and Prasad, 2019). This limitation has inspired an intense search for pro-immune modulators that could synergize with immune checkpoint blockade. sIL7R is one such pro-immune modulator and we propose its up-regulation could be used to enhance anti-cancer immunity and the response rates to immunotherapies. Supporting this premise, IL7 has been shown to augment the expansion of tumor-specific CD4^+^ (Ding et al., 2017) and CD8^+^ (Johnson et al., 2015) effector T cells, and to reduce tumor volume in diverse murine cancer models (Andersson et al., 2009; Ding et al., 2017; Li et al., 2007; Murphy et al., 1993; Pellegrini et al., 2009; Sharma et al., 2003), thereby implying that the IL7/IL7R pathway enhances T cell-mediated anti-cancer immunity. We predict that by potentiating IL7, sIL7R could improve the promising results observed with IL7. In support of this, co-administration of IL7 with chimeric sIL7R-Fc protein was shown to further enhanced T cell anti-cancer immunity, resulting in almost complete inhibition of tumor growth and increased survival in a murine model of lung cancer with accompanying increases in T cell activities (Andersson et al., 2011). Although the rationale behind this study was to test whether the IgG Fc portion could enhance the activity of antigen presenting cells, it is likely that sIL7R also contributed to the enhanced T cell activities and anti-tumor reactivity. Lastly, IL7 signaling in T cells decreases expression of the inhibitory receptor programed cell death 1 (PDCD1, PD1) (Boettler and von Herrath, 2012), and this effect is predicted to be enhanced by sIL7R, thereby potentially diminishing PD1-mediated T cell suppression. This is further supported by findings that the anti-tumor effects of *α*-PD1 and *α*-CTLA4 combination therapy require a functional IL7/IL7R pathway (Shi et al., 2016). Collectively, this body of work supports sIL7R as an attractive candidate to boost anti-cancer immunity, and positions pro-sIL7R ASOs IL7R-001 and IL7R-004 as potential drug candidates for cancer immunotherapy.

We conclude that the IL7R ASOs identified here are good candidates to control sIL7R expression for potential treatment of autoimmunity and cancers. Specifically, the anti-sIL7R ASOs IL7R-005 and IL7R-006 are promising candidates for treatment of MS and other autoimmune diseases where sIL7R is elevated (e.g., type I diabetes, rheumatoid arthritis and systemic lupus erythematosus), whereas the pro-sIL7R ASOs IL7R-001 and IL7R-004 are ideal candidates as novel immuno-oncology therapeutics to enhance anti-cancer immunity. Future preclinical studies will test the therapeutic potential of these anti-sIL7R and pro-sIL7R ASOs as viable candidates for treatment of autoimmunity and cancers, respectively.

## MATERIALS AND METHODS

### Generation of reporter plasmids and stable reporter cell lines

To create the GFP-IL7R reporter we subcloned the genomic sequence of IL7R previously used in the IL7R splicing reporter pI-11-IL7R (Gregory et al., 2007) into the pcDNA5/FRT/TO plasmid containing the coding sequence of the enhanced green fluorescent protein (eGFP) interrupted by the pI-12 intron (pGint) (Wagner et al., 2004). The region of IL7R spanned the last 614 bp of intron 5, the entire exon 6 and the first 573 bp of intron 6 and was cloned within the pI-12 intron of Gint using *XbaI* and *XhoI* restriction sites. The different alleles of the SNP rs6897932, as well as mutations to the 5’-splice site (5’Cons and 5’Mut) and exonic splicing elements (*Δ*ESE2 and *Δ*ESS3) in IL7R exon 6 were introduced using the QuikChange Lightning Site-Directed Mutagenesis kit (Agilent) following the manufacturer’s recommendations. The mutations were introduced as transversion substitutions and are shown in Fig. 1B. The pcDNA5/FRT/TO plasmid contains elements that enable generation of inducible expression (Tet-on) stable cell lines using the Flp-In T-Rex System (Thermo Fisher Scientific). Inducible HeLa stable cell lines expressing the different versions of the GFP-IL7R reporter (wild-type and mutants) were generated following the manufacturer’s protocol.

### Cells and cell culture

HeLa stable reporter cell lines were grown in DMEM medium (Thermo Fisher Scientific) supplemented with 10% heat-inactivated FBS free of tetracycline (Omega Scientific), 1% Penicillin-Streptomycin (Thermo Fisher Scientific), 2.5 μg/mL blasticidin (Invivogen), and 200 μg/mL hygromycin B (Thermo Fisher Scientific). Expression of the reporter was induced using 1 μg/mL doxycycline (Sigma-Aldrich). Primary CD4^+^ T cells from two healthy donors were purchased from Physician’s Plasma Alliance and cultured in RPMI 1640 medium (Thermo Fisher Scientific) supplemented with 10% heat-inactivated FBS (Omega Scientific) and 1% Penicillin-Streptomycin.

### Morpholino ASOs and transfection into HeLa cells and CD4^+^ T cells

Morpholino ASOs of interest were synthesized by Gene Tools. The control ASO (ASO-Ctrl) used is the standard control morpholino from Gene Tools. The sequence of these ASOs is shown in Table S1s. These ASOs were transfected at 10 μM into the HeLa reporter cell lines (5.0 × 10^4^ cells/well in 24-well plates) using 6 μM of the Endo-Porter PEG transfection reagent (Gene Tools) in the supplemented DMEM media. The cells were incubated with the corresponding ASO and Endo-Porter PEG for 36 hours, followed by induction of reporter expression by addition of 1 μg/mL doxycycline for 12 hours, at which time cells were collected for analysis of GFP expression and IL7R exon 6 splicing. The cells from each well were detached with 100 μL of Trypsin-EDTA 0.25% (Thermo Fisher Scientific), washed with PBS (Thermo Fisher Scientific) and the cell suspensions were split into two tubes, one for RNA isolation to quantify the effects on exon 6 splicing (by RT-PCR), and the other to quantify GFP expression (by Flow Cytometry). Dose-response studies were conducted similarly but ASOs were transfected at various concentrations (0, 1, 5 and 10 μM). All experiments using the reporter cell lines were conducted at least three times (each in triplicate wells) with similar results.

CD4^+^ T cells were transfected by nucleofection using a 4D-Nucleofector (Lonza). In brief, a total of 1.6 x 10^8^ cells were transfected with 2 μM of the corresponding ASO using the P3 Primary Cell 4D-Nucleofector X Kit L (Lonza) and the EO-115 program. The cells were incubated for 24 hours, washed with PBS, and 2.0 x 10^7^ cells from each cell suspension were plated per well in triplicate wells in 24-well plates. The cells were grown for 72 hours, at which time cells were collected for measurements of exon 6 splicing (by RT-PCR), sIL7R secretion (by ELISA using cell supernatants) and mIL7R cell surface expression (by Flow Cytometry). Experiments in CD4^+^ T cells were conducted three times (each in triplicate wells) with cells isolated from two healthy donors with similar results.

### RT-PCR analysis of IL7R exon 6 splicing

Total cell RNA was isolated from ASO-treated cells using the ReliaPrep RNA Cell Miniprep system (Promega) and treated in-column with DNase I to degrade DNA, following the manufacturer’s protocol. Either 1 μg (HeLa cells) or 0.1 μg (CD4 T cells) of total RNA was used as input for reverse transcription using the High-Capacity cDNA Reverse Transcription System (Thermo Fisher Scientific) following the manufacturer’s recommendations. Quantification of exon 6 splicing in transcripts from the reporter was conducted by PCR amplification with primers complementary to the coding sequence of GFP (Forward primer: 5’-CCACAAGTTCAGCGTGTCCG-3’; Reverse primer: 5’-CGTCCTTGAAGAAGATGGTGCG-3’), whereas for the endogenous IL7R, primers were complementary to IL7R exon 5 (Forward primer: 5’-GGCAGCAATGTATGAGATTAAAGTTCG-3’) and exon 7 (Reverse primer: 5’-CAAAGATGTTCCAGAGTCTTCTTATGATC-3’). PCR reactions were prepared as follows: 5 μL of the corresponding RT reaction (diluted 1:5) was mixed with 200 nM of the corresponding forward and reverse primers, 100 nM dNTPs, 50 mM KCl, 2 mM MgCl_2_, and 0.4 μL of Taq polymerase. PCR products were resolved and quantified by molarity on a 2100 Bioanalyzer Instrument (Agilent) using the High Sensitivity DNA Kit (Agilent). Percent of exon 6 exclusion was determined as: [% exon 6 exclusion = excluded product / (excluded + included products) x 100%]. In each figure, the data are presented as the mean of triplicate wells from a representative experiment and error bars represent standard deviation.

### Quantification of GFP by flow cytometry

To quantify GFP expression from the various versions of the GFP-IL7R reporter (wild-type and mutants), the reporter cell lines were resuspended in PBS at 500 cells/μl, and 100 μl of each cell suspensions were loaded per well in round-bottom 96-well plates (Fisher Scientific). The samples were assayed in a Guava EasyCyte 6-2L HT Flow Cytometer (Luminex), and GFP expression data was extracted as Mean Fluorescence Intensity (MFI) in the Green 525/30 channel. MFI values were converted to Log2(fold-change) over control (ASO-Ctrl) to enable direct comparison of ASOs that increase or decrease GFP expression. For the experiments comparing the C-allele versus the T-allele of the SNP rs6897932 (Figs. 4 and 5), the data is shown as MFI relative to the T allele reporter cell line treated with ASO-Ctrl (Fig. 4) or relative to the C allele reporter cell line treated with ASO-Ctrl (Fig. 5).

### Quantification of sIL7R by ELISA

Supernatants from ASO-treated HeLa cells or human primary CD4^+^ T cells were collected and 400 μL of each were concentrated to 100 μL by centrifugation using Amicon Ultra 3K centrifugal filters (Millipore Sigma), and the 100 μL of the concentrated supernatants were used to quantify sIL7R secretion in ELISA assays. Measurements of secreted sIL7R in HeLa cell supernatants were conducted as follows: 96-well plate (R&D Systems) were coated at 4^°^C overnight with a mouse anti-human IL7R monoclonal antibody (R&D Systems, # MAB306). The next day, the plates were washed with PBS and blocked with 3% BSA in PBS for one hour at room temperature, followed by two-hour incubation at room temperature with the concentrated supernatants, and four washes with PBS-Tween (0.05%). Detection of bound sIL7R was carried out by one-hour incubation at room temperature with biotinylated goat anti-human IL7R polyclonal antibody (R&D Systems, # BAF306), followed by 30-minute incubation at room temperature with streptavidin-horseradish peroxidase (Millipore Sigma, # 18-152), and 20-minute incubation at room temperature with TMB peroxidase substrate (SurModics BioFX), with four washes with PBS-Tween (0.05%) between manipulations. The reaction was stopped with 1N sulfuric acid (H_2_SO_4_), and the product was immediately visualized in a plate reader at 450 nm. The concentration of samples was extrapolated from standard curves of recombinant human IL7R-Fc chimera (R&D Systems, # 306-IR).

Quantification of secreted sIL7R in CD4^+^ T cell supernatants was conducted with the IL7R (CD127) ELISA kit (My BioSource, # MBS824893), following the manufacturer’s protocol. In brief, 100 μl of concentrated supernatants were pipetted into a 96-well plate coated with an immobilized anti-human CD127 antibody. After a 90-minute incubation at room temperature the plate was washed three times in wash buffer and a biotin-labeled detection antibody was added to detect the bound sIL7R. The plate was incubated for 60 minutes at 37°C, washed three times with wash buffer, followed by a 45-minute incubation at 37°C with a streptavidin-horseradish peroxidase, five washes with wash buffer, and a 30-minute incubation at 37°C in the dark with TMB peroxidase substrate. The reaction was stopped with 2N sulfuric acid (H_2_SO_4_) and the absorbance was immediately read at 450nm in a microplate reader. The concentration of supernatant samples was extrapolated from standard curves of recombinant human CD127 protein supplied in the kit.

### Quantification of mIL7R by flow cytometry

Cell surface expression of mIL7R was quantified in CD4^+^ T cells by staining of mIL7R in intact, live cells. In brief, 1.0 x 10^6^ of the ASO-treated cells were resuspended in 250 μL of PBS supplemented with 5% FBS and incubated on ice for 30 minutes to block Fc receptors. The cells were then centrifuged at 250 x g for 5 minutes at 4^°^C, the PBS was removed, and the cells were resuspended in 200 μL of PBS supplemented with 1% FBS and containing 2.5 μL of Alexa Fluor 647-conjugated anti-human CD127antibody (BioLegend) or Alexa Fluor 647-conjugated mouse IgG1 k isotype control (BioLegend). The cells were incubated with the corresponding antibody at room temperature for one hour, washed three times with 500 μL PBS with 1% FBS, resuspended in 200 μL PBS, and loaded into a well of a round-bottom 96-well plate. Data on mIL7R cell surface expression was extracted as Mean Fluorescence Intensity (MFI) in the Red-R 661/15 channel of a Guava EasyCyte 6-2L HT Flow Cytometer. MFI values are shown in Figure 3C relative to ASO-Ctrl.

### Statistical assessment

In all experiments, statistical significance was assessed by two-sided Student’s t test pairwise comparisons against ASO-Ctrl, except for dose-response experiments were comparison was against 0 μM (* *p < 0.05*; ** *p < 0.01*; *** *p < 0.001*).

## SUPPLEMENTAL MATERIAL

Supplemental material, including supplemental figures and tables, is available for this article.

## ACKNOWLEDGEMENTS

We thank Alexandra Vincent and Jon D. Moulton from Gene Tools for their technical advice in the design of morpholino ASOs. This study was funded by startup funds from the University of Texas Medical Branch (M.A.G.-B.), a Technology Commercialization Plan Grant from the University of Texas Medical Branch (G.G.-M. and M.A.G.-B.), and NIH Grants R41-AI141323 and R41-AI149920 to Autoimmunity BioSolutions (G.G.-M.). The University of Texas has filed patent applications covering the ASOs described here, where M.A.G.-B., G.G.-M. and S.B.B. are inventors. M.A.G.-B. and G.G.-M. co-founded Autoimmunity BioSolutions, a biotech company developing these ASOs for treatment of autoimmune diseases and cancers. S.B.B. is a consultant for Autoimmunity BioSolutions. They attest that in no way did these interests influence the planning, execution or interpretation of the results, or the writing of this manuscript. The other authors have no competing interests or other interests that might be perceived to influence the results and/or discussion reported in this paper.

